# A biomechanical comparison of the jerk, back jerk, vertical jump and jump squat

**DOI:** 10.1101/2021.03.15.435446

**Authors:** Jiatian Liu

**Author notes:** (JL).

## Abstract

Although weightlifting exercises and their pulling and catching derivatives have been well studied, less is known about the group of weightlifting overhead pressing derivatives (WOPDs). There were two purposes of the present study. Above all, it was to compare WOPDs (i.e. jerk and back jerk) with jump squat in enhancing sport performance such as vertical jump in the perspective of biomechanics. Furthermore, it was to gain a better understanding of the biomechanical difference between jerk and back jerk. The study compared the kinetics, kinematics, and muscle activation of the jerk, back jerk, vertical jump and jump squat. Ground reaction forces, joint angle data and electromyography were collected from 20 track and field athletes while they performed the four movements. The joint coordination pattern and EMG characters of the jump squat were more similar to those of the vertical jump. WOPDs, especially jerk, exhibited less similarity with vertical jump in joint coordination, as well as the subsequent peak activation. The electromyography data demonstrated significant differences in rectus femoris and gluteus maximus in the relative timing of peak activations and the maximum activation. Jump squat was greater in peak force (2211N) than jerk (P < .005), greater peak power (4749W) than jerk and back jerk (P < .001) and no significant difference with WOPDs in rate of force development. Back jerk produced a greater peak force (2223 N), peak power (3767W) and eccentric RFD (15965 N·s^−1^) than jerk (2061N)(P < .001) (3364W)(P < .05) (12280 N·s^−1^)(P < .01)and the opposite in concentric RFD that jerk (7702 N·s^−1^) was greater than back jerk (5972 N·s^−1^)(P < .05). We can conclude that for runner and jumpers, jump squat was better for improving vertical jump than WOPDs and back jerk is better than jerk in power training.

## Introduction

Weightlifting derivatives, which consisted of weightlifting catching, pulling and pressing derivatives [1–3], has been applied to enhance sports performance such as sprinting and jumping [4, 5]. Although the pulling and catching derivatives have been extensively studied [6–8], there were far less researches about the group of weightlifting overhead pressing derivatives (WOPDs).

One of the major exercises of WOPDs is the jerk, which is not technically a pressing motion. Instead, the propulsion phase was that the bar was accelerated vertically via triple extension (hips, knees and ankles) of the athlete, then quickly transferred into the catch position that the athlete dropped underneath the bar. The WOPDs such as jerk, push press, split jerk or back jerk has been largely implemented in strength and conditioning programs of a wide range of sport population for improving performance [9–12]. Previous literatures suggested that these exercises are powerful tools for development of strength and power [13–15], as well as that the fast triple extension of lower body resembled many sporting activities [16].

The mechanical similarity between a training exercise and the target sport performance was a foremost prerequisite for selection of exercises to improve physical ability. It also resulted in the classification from general to specific in respect to relationships to a specific sport movement. After comparison of kinetic and kinematic qualities, selection of the most identical exercises will contribute to a more direct transfer from the training exercises to sporting tasks [17].

In the limited researches of WOPDs, the mechanical similarity were reported between WOPDs and specific sport movements. Cleather et al. [18] conducted a study on vertical jump and jerk by calculating the intersegmental moments expressed at hip, knee and ankle joints. The main finding was that jerk was more like a general training modality than specific in relation to jumping. However, Cushion et al. [19] indicated that push jerk was more similar to countermovement jump (CMJ) than jump squat with the notion that jump squat is one of the commonly implemented exercises to strengthen jumping abilities. In contrast, other researchers, who also compared jerk, jump squat and CMJ, suggested that the loaded jump squat exercise was more related to jumping and sprinting abilities than to the push press [20, 21]. Although motions of lower body were all triple extensions, the nature of movements in these exercises was different. The jumping movements were about to overcome the inertia of the whole system (body and external load), rather the jerking movements focused on transfer of energy from the body to the external load.

The discrepancy from the different results of previous studies was yet to be resolved, which provoked a thought that whether the coordination between adjacent joint was different with these exercises. And no previous studies have assessed the kinematics in the perspective of joint coordination. The continuous measure of joint coupling are vector coding technique and continuous relative phase (CRP). Due to the limitation that CRP may produce inaccurate results in non-sinusoidal time series of joint motion. Quantification of joint coordination was obtained by a modification of a vector coding technique suggested [22]. Using the relative motion plots, it showed the synchrony between the motions of two joints. The discrepancy and lack of joint coordination from previous literatures highlighted the importance of studying WOPDs and their correspondence to specific motor quality one stage further and therefore, also sport performance.

In addition, features such as the initial position of the barbell and the style of catching resulted different power output values. Flores et al. [23] compared the distinctions between the jerk from the chest and the jerk from the back throughout different relative intensities. The results indicated that across all the loads assessed greater power output was achieved by the jerk from the back other than the jerk from the chest. And both exercises elicited the peak power at a relative intensity of 90% of 1RM, while greater for the jerk from the back than the jerk from the chest (3400.22 ± 691.07 and 3103.34 ± 616.87 W, respectively), but not significantly. Considering the power was calculated without the lifter’s body weight and all the subjects were professional weightlifters, some questions, such as whether subjects from other sports (e.g., sprinters, jumpers, or throwers) would elicit different results and how the kinematic differences observed between jerk and back jerk may affect the kinetic values, were still unclear. These would provide helpful knowledge for athletes and coaches when they choose training exercises so as to improve performances.

Therefore, the aim of this study was to compare WOPDs (jerk and back jerk) and jump squat and their advantages in enhancing sport performance such as vertical jump by means of kinetics, kinematics and muscle activation. A secondary aim was to gain a better understanding of the biomechanical difference between jerk and back jerk. It was hypothesized that back jerk is better than jerk in developing force and power, but nowhere near jump squat in similarity of CMJ.

## Materials and methods

### Participants

Twenty university aged male track and field athletes (age: 21.05±2.09 years; height: 176±5.10 cm; mass: 73±8.33 kg; squat 1RM: 107.05±16.03 kg) volunteered for the study and provided written consent including experimental procedures and risks before participation. This study was approved by the institutional review board of Beijing Sports University and conformed to the principles of the World Medical Association’s Declaration of Helsinki. All the participants were free from injury with a minimum of 3 years performing each exercises and were asked to avoid competitive sport at the time of testing.

### Procedures

1 repetition maximum (1RM) test was conducted at least 1 week before the main testing session. All participants were required to perform a 1RM back squat based on the testing protocol of Winchester et al. [24].

The load used in the jump squat was 40% of 1RM back squat, which was suggested as a typical training load in literatures [25]. The same weight was used for both jerk trials because the barbell weight associated with 40% 1RM back squat was well within the range of loads previously suggested in the literature for jerk training [26]. Moreover, the purpose of the study was to compare movements of each training exercises, it is necessary to reduce the impact from other variables such as load. In view of this, the same absolute and typical load used for all three loaded lifts was a more rigorous method for comparison of movements.

The main testing session began with a standardized warm-up consisting of 10 bodyweight squats, 10 inchworms and barbell work including 10 jumps squats, 10 jerks and 10 back jerks. And during the 10 loaded lifts, the load increased from 20kg to their tested load in their own pace. Participants then performed four exercises in a randomized order. Subjects performed two successful trials for each exercise with self-selected rest periods between each repetition. Previous research has established that trained subjects show a high degree of reliability between repetitions when self-regulating rest periods [27]. Because athletes were experienced in training in the present study, they were trusted in their judgment to self-select rest periods; this was also to ensure that they felt adequately recovered between each repetition.

### Countermovement Jump

The CMJ began with subjects in an upright position with their arms free to swing during the movement. Subjects were instructed to jump maximally for each repetition with depth of the countermovement jump self-selected.

#### Jump Squat

The loaded JS began with subjects in an upright position with the barbell placed on the upper back. Subjects performed a maximal jump initiated with a countermovement where depth was again self-selected.

#### Jerk

Jerk is called front jerk (FJ) for differentiation in current study. The loaded FJ began with subjects in an upright position with the barbell placed on the anterior deltoids. Subjects initiated the movement with a countermovement before extending the arms above the head and landing in a semi-squat position.

#### Back Jerk

The loaded BJ began with subjects in an upright position with the barbell placed on the shoulders and upper back (as in the jump squat position). Subjects initiated the movement with a countermovement before extending the arms above the head and landing in a semi-squat position.

GRF data were collected on a separate piezoelectric force plate (Kistler 9281CA, Switzerland) under each foot, in a capture area defined by an 8-camera motion analysis system (Motion Analysis Raptor-4, USA). Markers were placed on bony landmarks of anatomical structures on the shoulder (acromioclavicular joint), hip (greater trochanter), knee (lateral ridge of tibial plateau), ankle (apex of the lateral malleolus), and distal end of the foot (metatarsus head). Surface electrodes (Trigno Wireless EMG,DELSYSI.,Natick,USA) were fixed to the belly of the medial gastrocnemius, vastus medialis, rectus femoris, biceps femoris, and gluteus maximus, parallel to the muscle fibers on the dominant leg.

### Data collection and analysis

The motion analysis system and force plate data was sampled at 200 Hz and 1000 Hz and synchronized with each other by saving the force plate data into the motion analysis system. The commencement of upward phase was the moment when the hip, knee, or ankle joint angle began to increase (extend) after the countermovement phase. The completion of the upward phase was the moment when extension of all joints were achieved.

Joint coordination was evaluated by a modification of a vector coding technique suggested [28]. Angle–angle plots were constructed for coupling of two adjacent joints with distal motion plotted on the ordinate and proximal motion plotted on the abscissa during the upward phase of triple extensions. The absolute resultant vector between two adjacent data points were calculated (Eq. (1)). Thus, coupling angle was the angle between the absolute resultant vector and its orientation to the right horizontal. Following conversion from radians to degrees, the resulting range of values for Φ, or coupling angle, was 0–90°.

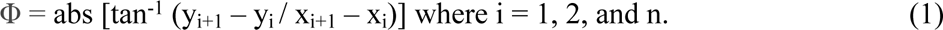

Each exercise has two coupling angles which are hip-knee (HK) and knee-ankle (KA) coupling angles. A coupling angle of 45° would indicate equal amounts of distal and proximal joints motion. An angle greater than 45° indicates greater distal motion relative to proximal motion. A mean coupling angle was calculated for each of the subjects for the upward phase.

Peak force, peak rate of force development (instantaneous), peak power as well as the percentage of upward movement when these peak values reached were calculated by analyzing data from force plate. At the beginning of each exercise, the mass of the system was measured by participants stood motionless on the force plate and calculate the net vertical force acting on the system. Calculation of the instantaneous vertical velocity of the system was to integrate the net vertical force with respect to time, and divide by mass. Instantaneous power was calculated as the product of the instantaneous vertical velocity and instantaneous vertical ground reaction force.

Surface electromyography (EMG) was sampled at 2000 Hz and was synchronized with the motion analysis data. Using delsys EMGworks4.0 Analysis, EMG data were analyzed by using a band-pass, 4th order Butterworth filter with a 10-400Hz cut-off. Then rectified and filtered using a 4th order, low pass Butterworth filter with a cut-off frequency of 20 Hz. Finally, linear enveloped curves was achieved by using a customized MatLab (MatLab R20016b, Math Works, Natick, MA, USA) script. The signal that greater than 20% of the maximum signal attained during the trial was recognized as muscle activation.

In order to express joint angular displacement as percentage of the upward phase, each joint angular displacement signal of the upward phase was interpolated by Origin (Origin Pro9, Originlab, Northampton, MA, USA) to generate 100 data points. The same procedure was applied to the force plate and EMG data as standardization of all data, which facilitated statistical comparison. It also allowed data of all participants to be graphed concurrently as the mean ± 95% confidence interval.

### Statistical analysis

The vertical jump and jump squat trials with the highest jump height, and the two jerks with the highest bar height prior to the catch, were selected for statistical analysis. Differences in all variables measured during the trials were analyzed using one-way repeated measures analyses of variance (ANOVAs). The independent variable (exercise) of within participants had four levels: jump squat, jump, back jerk and front jerk. If the assumption of sphericity was not met, as determined using Mauchly’s Test, then Greenhouse-Giesser corrections were applied. When significant value was determined, Bonferroni post-hoc tests were used to determine where significant differences exited between exercises with adjustments to control for Type I error. Effect sizes were estimated using partial eta squared (η^2^_p_). Statistical analyses were performed using SPSS 19.0 (SPSS Inc, Chicago, IL, USA). Significance for all statistical tests was defined as P ≤ .05.

## Results

In HK coupling, coupling angle of jerk (56°) and back jerk (51°) were more knee dominated than jump squat (49°) and vertical jump (47°), while jerk was significantly different with all other three exercises and back jerk was significantly different with vertical jump (F(3, 57) = 24.590, P<.001, η^2^_p_ = .564) (Fig 1A). In KA coupling, mean coupling angle of jerk (32°), back jerk (29°), jump squat (29°) and vertical jump (27°) showed motion of knee were leading in more times relative to ankle, while jerk was significantly different with jump squat and back jerk (F(3, 57) = 4.849, P<.005, η^2^_p_ = .203) (Fig 1B).

**Fig 1.**
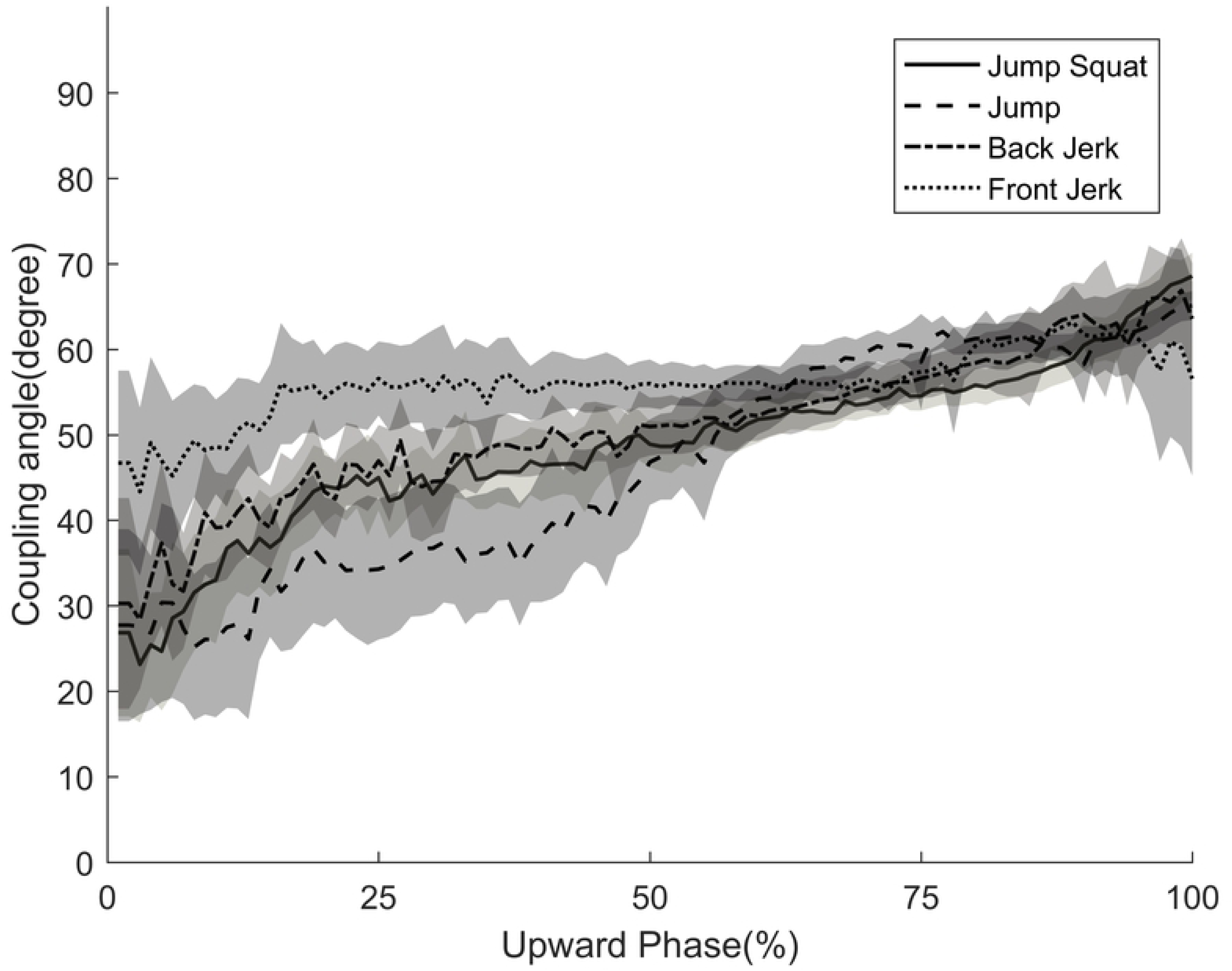
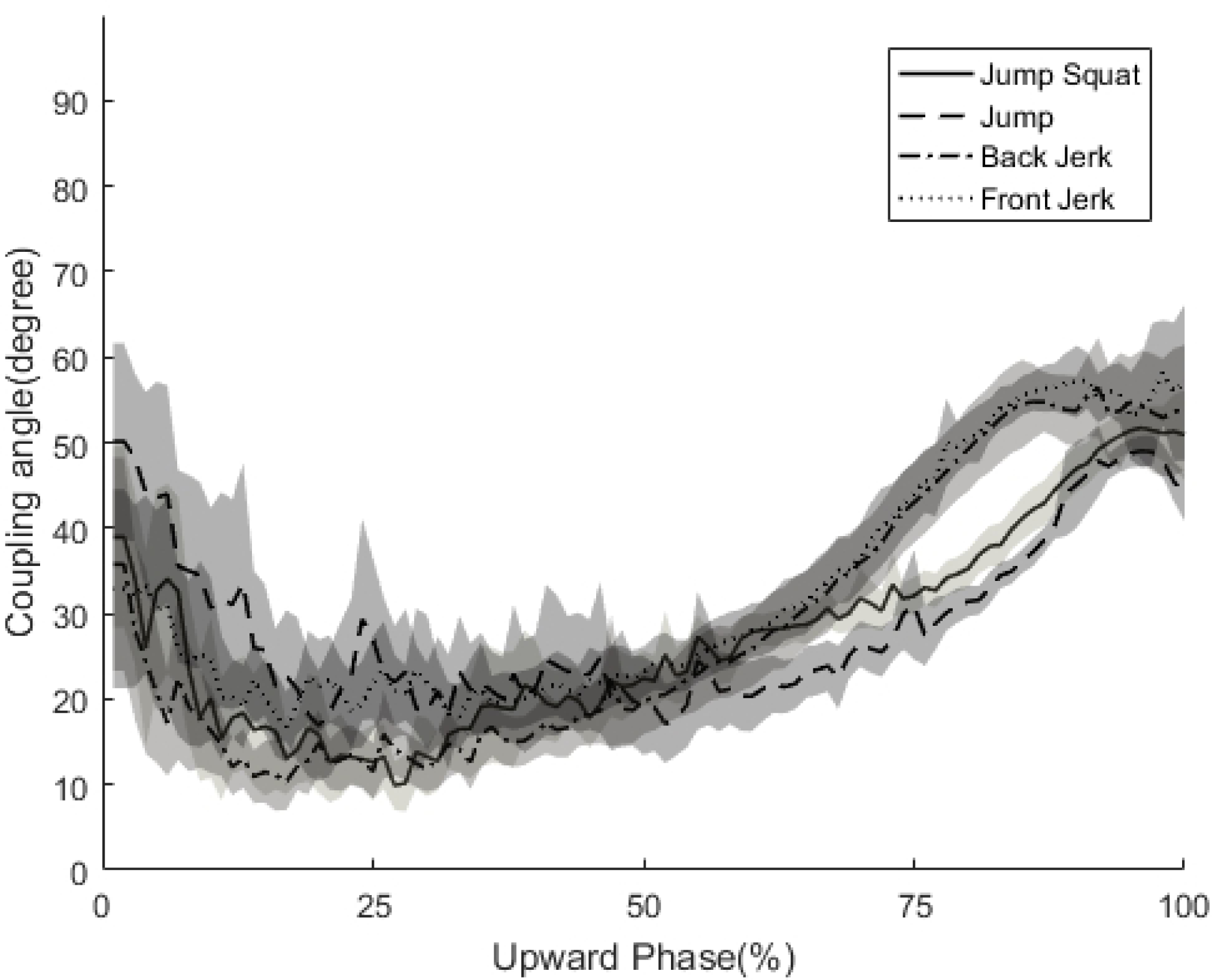
Upward phase of the (A) hip-knee coupling angle and (B) knee-ankle coupling angle during the upward phase of the jump squat, jump, back jerk and front jerk. Curves are ensemble average for all participants, while the shaded bands represent the 95% confidence interval.

On average, back jerk (2223N) and jump squat (2211N) elicited a significantly greater peak force than front jerk (2061N) and vertical jump (1923N), while front jerk elicited a significantly greater peak force than vertical jump (F(2.119, 40.259) = 28.868, P<.001, η^2^_p_ = .86)(Fig 2A). On average, the greatest peak power was observed in the vertical jump condition (6373W) and was significantly greater than the peak power generated in the jump squat (4749W), back jerk (3767W), and front jerks (3364W), which were significantly different from each other (F(1.655, 31.437) = 99.536, P<.001, η^2^_p_ = .84)(Fig 2B). On average, the greatest rate of force development was generated during the vertical jumps (10175 N·s^−1^), then the jump squat (7177 N·s^−1^) and back jerk (5972 N·s^−1^), which was significantly greater from each other, except that the front jerk (7702 N·s^−1^) was only significantly greater than the back jerk (F(1.748, 33.215) = 11.060, P<.001, η^2^_p_ = .368)(Fig 2C). The reliability of the rate of force development measure was estimated by calculating Cronbach’s alpha for the two trails. The value were .872, .974, .878 and .801 for jump squat, jump, back jerk and front jerk.

**Fig 2.**
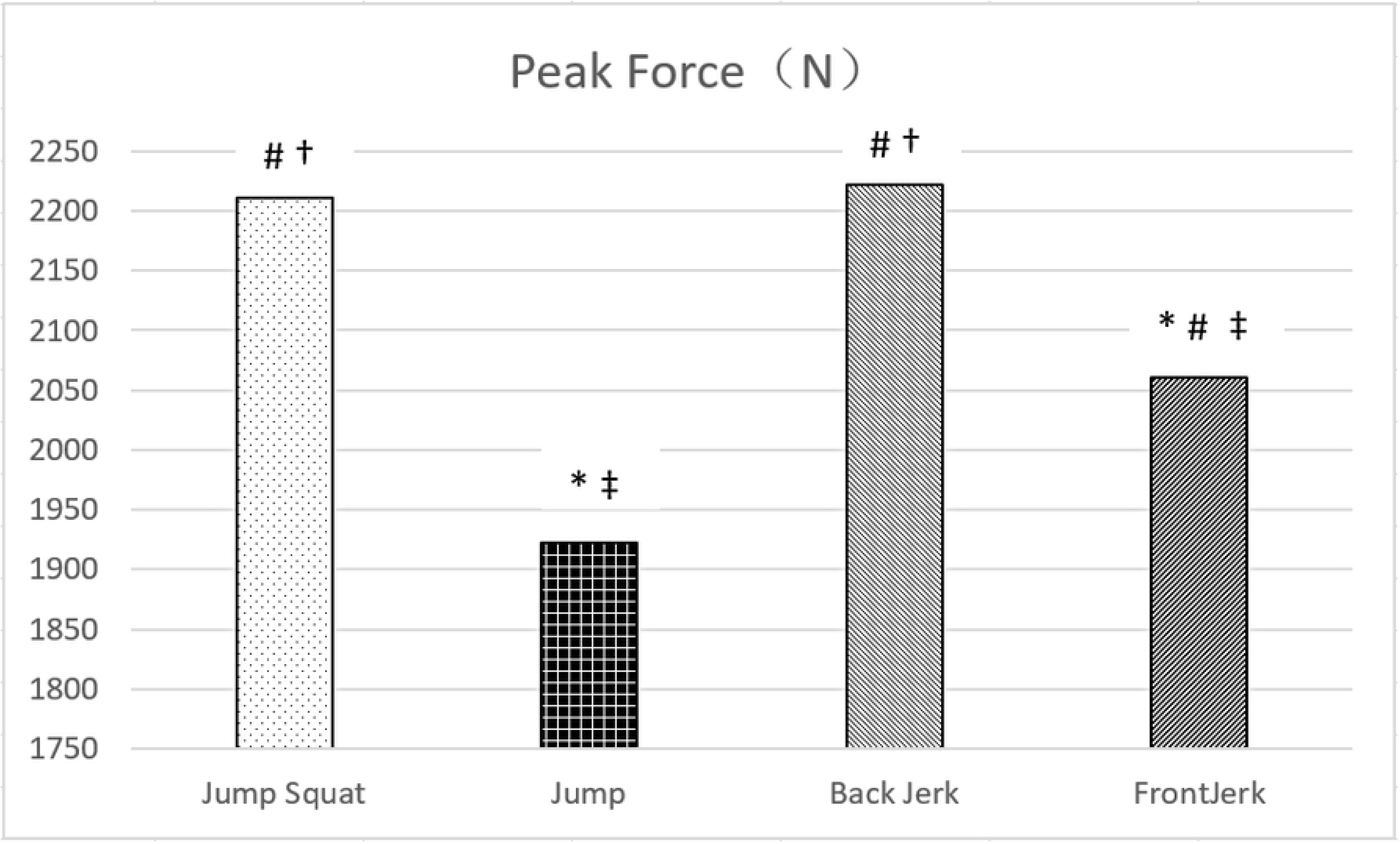
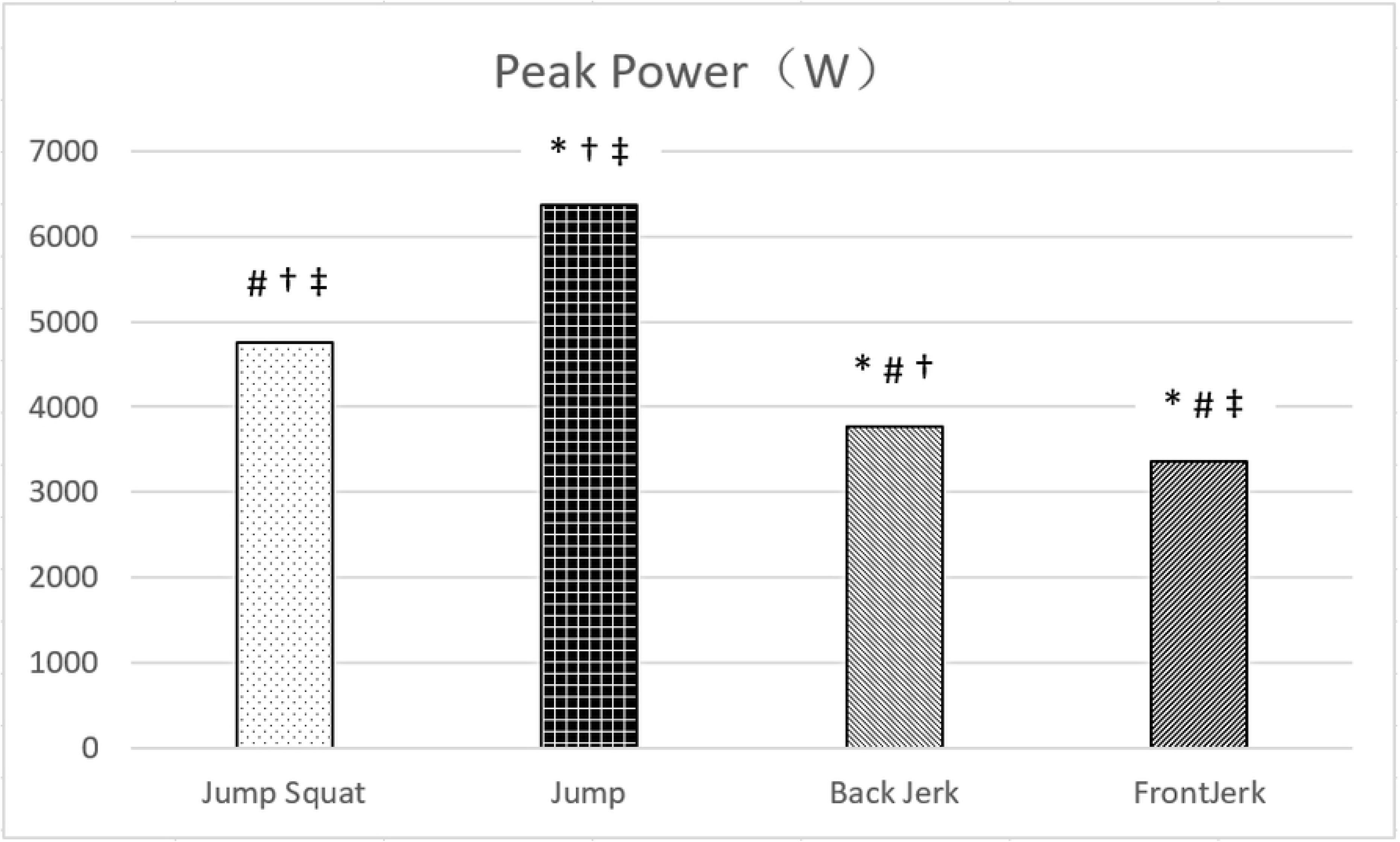
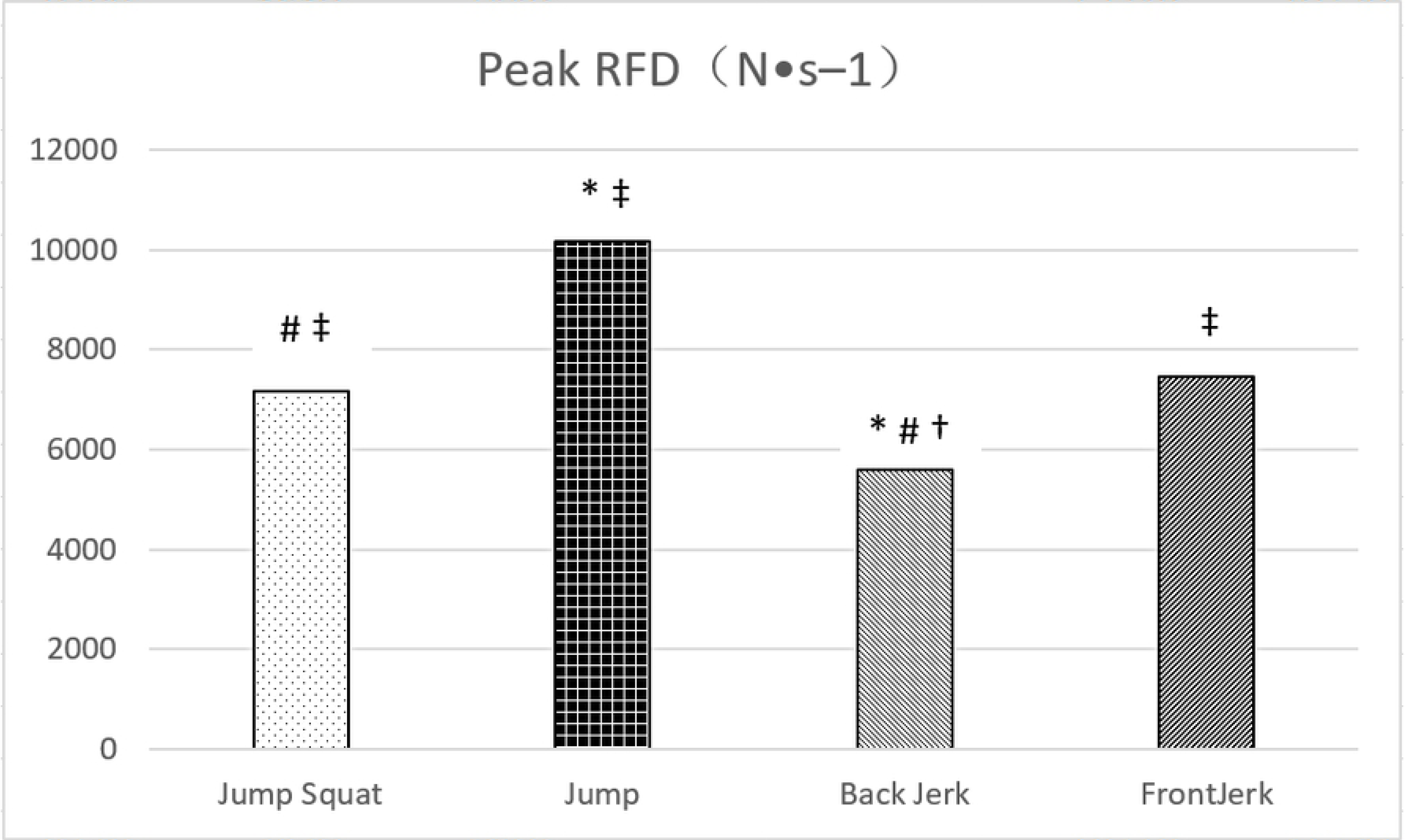
(A) Peak vertical ground reaction force, (B) peak power generated and (C) peak rate of force development (RFD) during the upward phase of the jump squat, jump, back jerk and front jerk. #indicates significantly different from vertical jump (P ≤ .05). *indicates significantly different from jump squat (P ≤ .05). †indicates significantly different from front jerk (P ≤ .05). ‡indicates significantly different from back jerk (P ≤ .05)

On average, peak force was achieved significantly earlier in the back jerk (18%) as a percentage of upward movement than the front jerk (41%), vertical jump (46%) and jump squat (56%), which were not significantly different from each other during the upward phase (F(3, 57) = 8.470, P < .001, η^2^_p_ = .308)(Fig 3A). Peak power was achieved first during the back jerk (57%) and front jerk (62%) which was not significantly different between each other, then followed by jump squat (75%) and vertical jump (78%), which was not significantly different between each other. Only both jerks were significantly greater than both jump squat and vertical jump (F(3, 57) = 54.552, P < .001, η^2^_p_ = .742)(Fig 3B). Peak rate of force development was achieved significantly earlier during the front jerk (30%) than the back jerk (34%), jump squat (47%) and vertical jump (48%), which only front jerk was significantly earlier than vertical jump (F(3, 57) = 3.809, P < .05, η^2^_p_ = .167).

**Fig 3.**
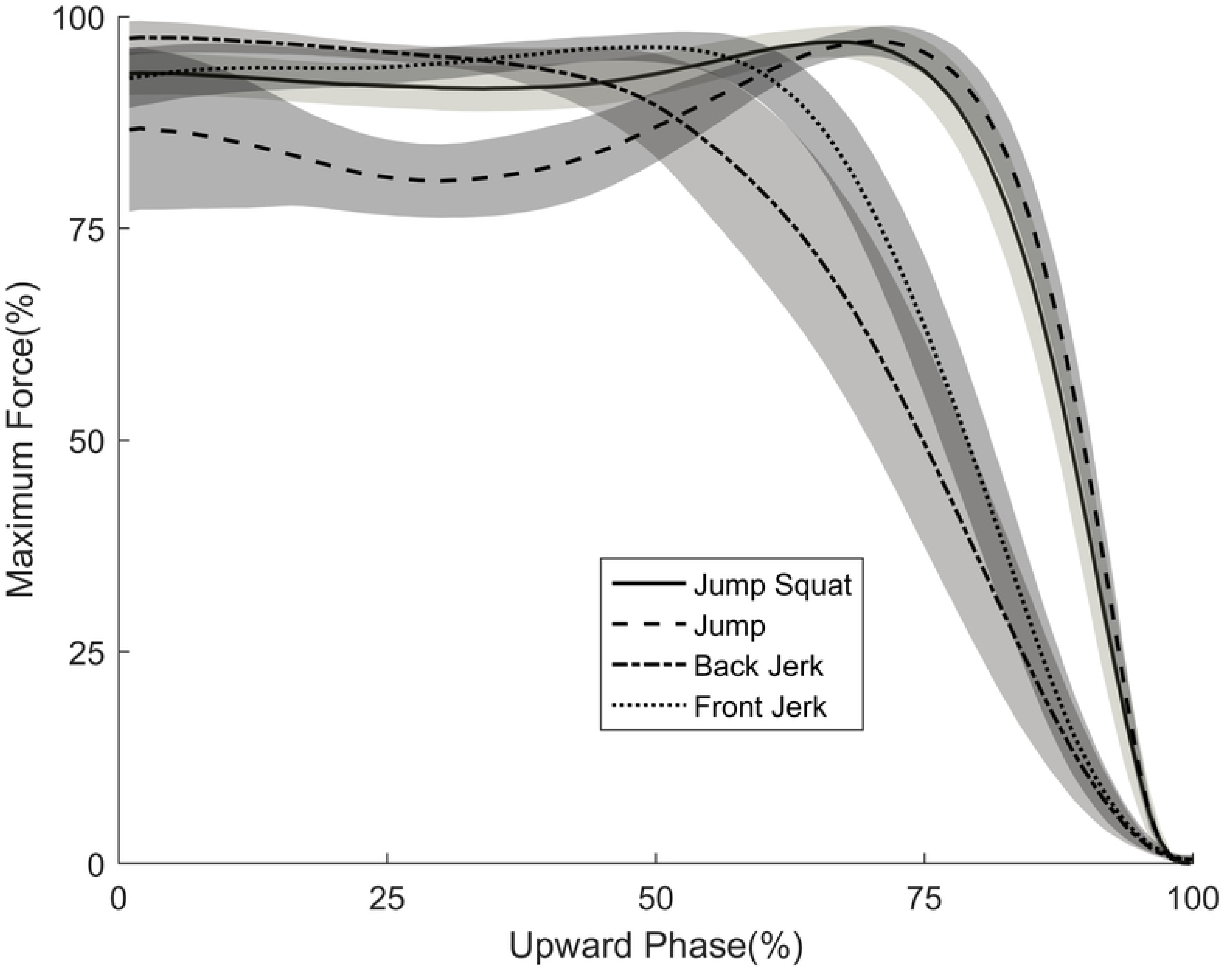
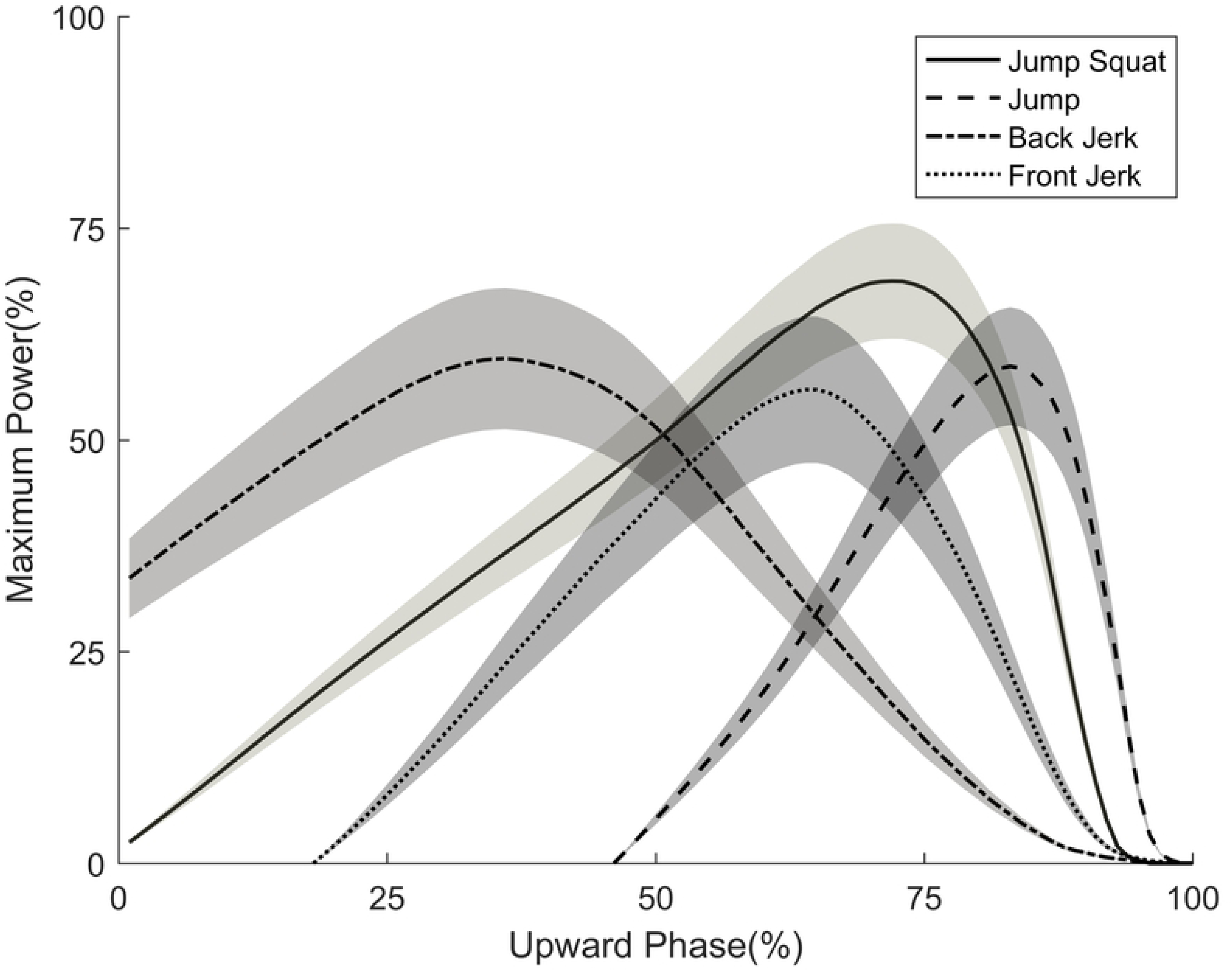
(A) Vertical ground reaction forces and (B) power as a percentage of the maximum values attained during the upward phase of the jump squat, jump, back jerk and front jerk. Curves are ensemble averages for all participants, while the shaded bands represent the 95% confidence interval.

The absolute timing of exercises are impossible to observe by representing dependent variables as a function of percentage of events, although it facilitates a statistical comparison. Such as the absolute timing of peak force is demonstrated by examination of vertical ground reaction force curves, which offered a qualitative suggestion of the timing of peak rate of force development (Fig 4A). Fig4B provides representative power curves of one participant whose body weight was 83 kg and 54kg of barbell mass used during the trials. Though this paper mainly focused on upward phases of the exercise, some findings during countermovement phase were worthy of note. Rates of force development were found significantly higher than concentric phase in jump squats (t(19) = 5.989, P<0.001), vertical jumps (t(19) = 5.940, P<0.001), back jerks (t(19) = 6.973, P<0.001) and front jerks (t(19) = 6.343, P<0.001). These findings can be discovered qualitatively by observing the slope in Fig 4A. During the countermovement phase, the greatest rates of force development was achieved in vertical jump (18044 N·s^−1^), then back jerk (15965 N·s^−1^) and jump squat (15039 N·s^−1^), which were not significantly different from each other. Except front jerk (12280 N·s^−1^) was significantly lower than the other three exercises (F(3, 57) = 11.253, P<.001, η^2^_p_ = .372).

**Fig 4.**
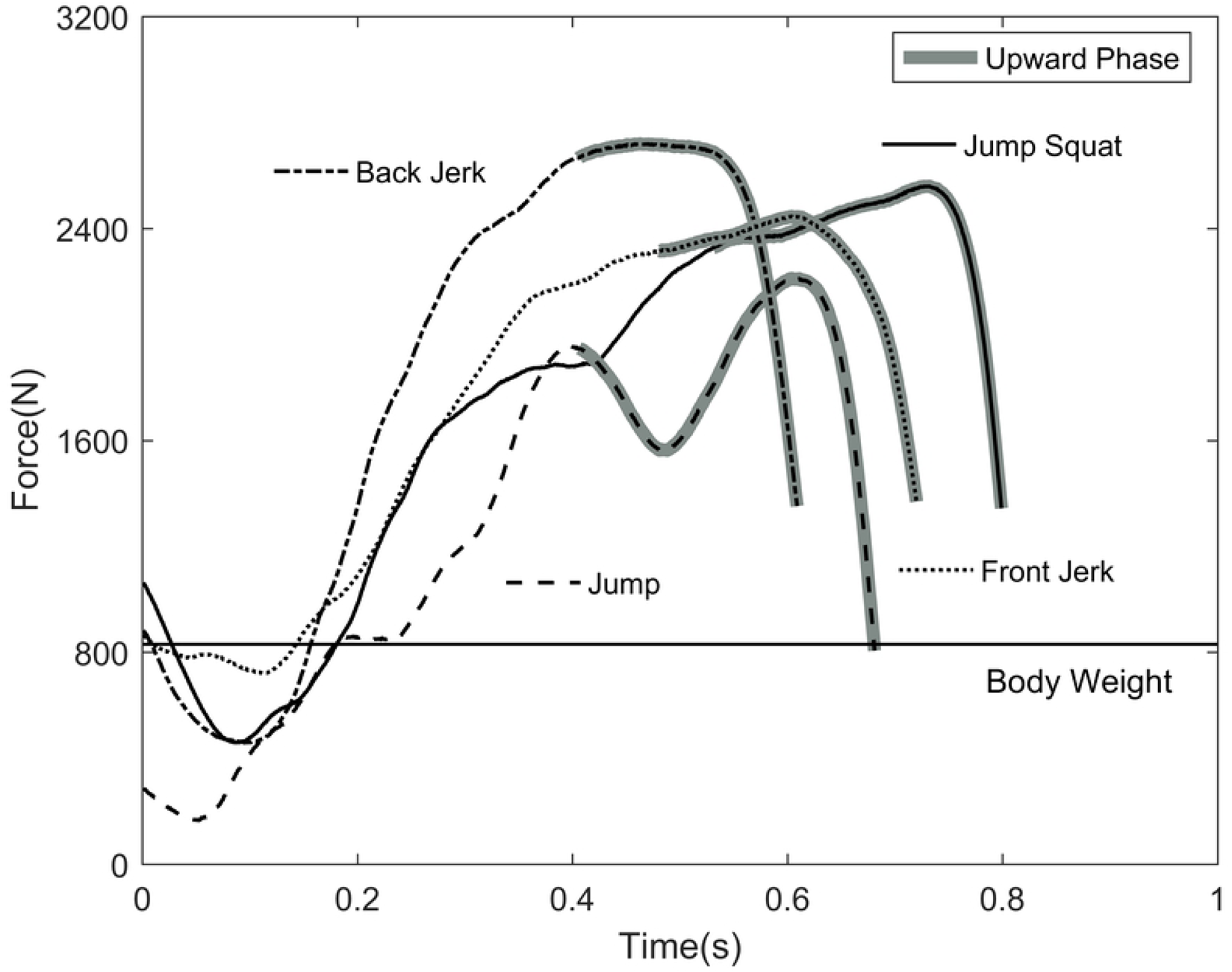
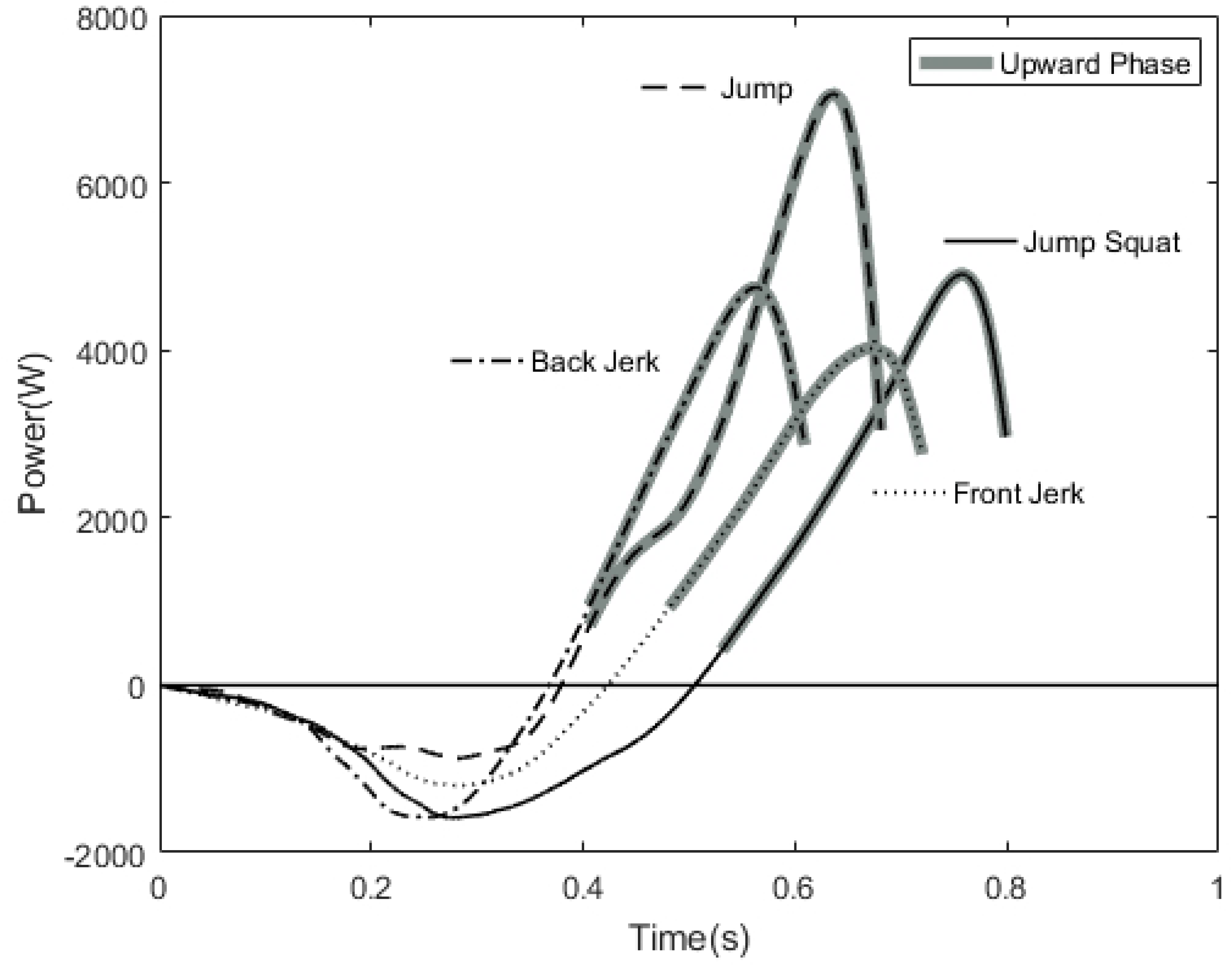
(A) Typical vertical ground reaction force and (B) power profiles, for Participant 7, for the jump squat, jump, back jerk and front jerk. Curves start at the initiation of the downward phase, while the shaded portions indicate the upward phase. Curves end when the weight of the participant, or participant and barbell, exceeded the magnitude of the vertical ground reaction force.

On average, the earliest peak EMG value of rectus femoris occurred was front jerk (37%), then back jerk (45%), vertical jump (66%) and jump squat (72%), while front jerk was significantly earlier than vertical jump and jump squat and back jerk was only significantly earlier than jump squat (F(3, 57) = 7.069, P<.001, η^2^_p_ = .271) (Fig5A). In vastus medialis, order of peak EMG value was similar that front jerk (32%) is earliest, then back jerk (33%), jump squat (45%) and vertical jump (56%), which were not significantly different between each other (F(3, 57) = 3.384, P<.05, η^2^_p_ = .271) (Fig5B). On average, peak EMG value of gluteus maximus was earlier in vertical jump (27%), then jump squat (31%), back jerk (34%) and front jerk (47%), while only vertical jump was significantly earlier than front jerks (F(3, 57) = 3.274, P<.05, η^2^_p_ = .174) (Fig5C). On average, peak EMG value of biceps femoris was earlier in back jerk (50%), then vertical jump (54%), front jerk (61%) and jump squat (75%), while only back jerk was significantly earlier than jump squat (F(3, 57) = 4.685, P<.01, η^2^_p_ = .198)(Fig5D). On average, peak EMG value of medial gastrocnemius was earlier in front jerk (56%), then back jerk (59%), vertical jump (71%) and jump squat (76%), while only jump squat was significantly later than both jerks and no significant difference between two jerk conditions (F(3, 57) = 6.559, P<.005, η^2^_p_ = .257) (Fig5E).

**Fig 5.**
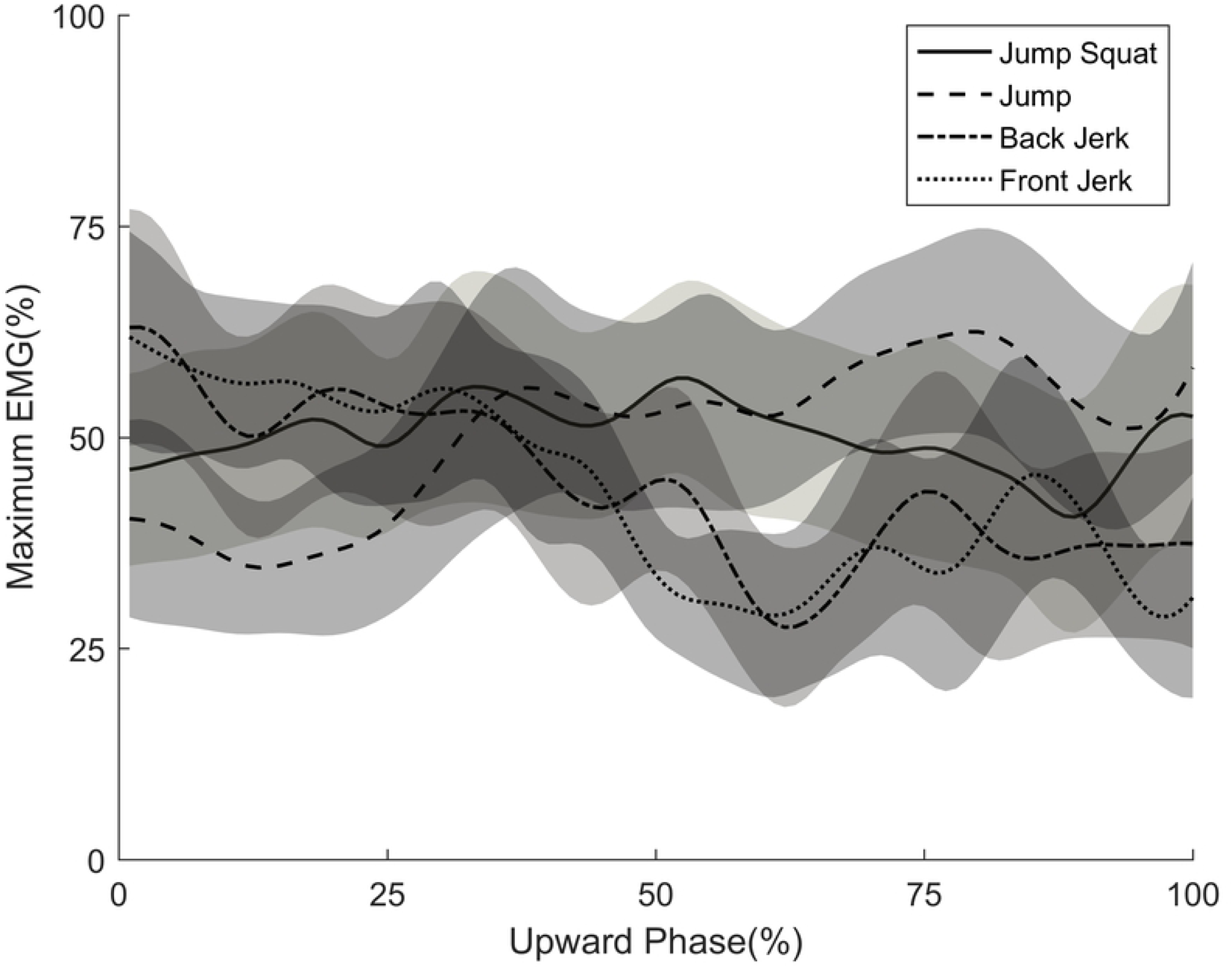
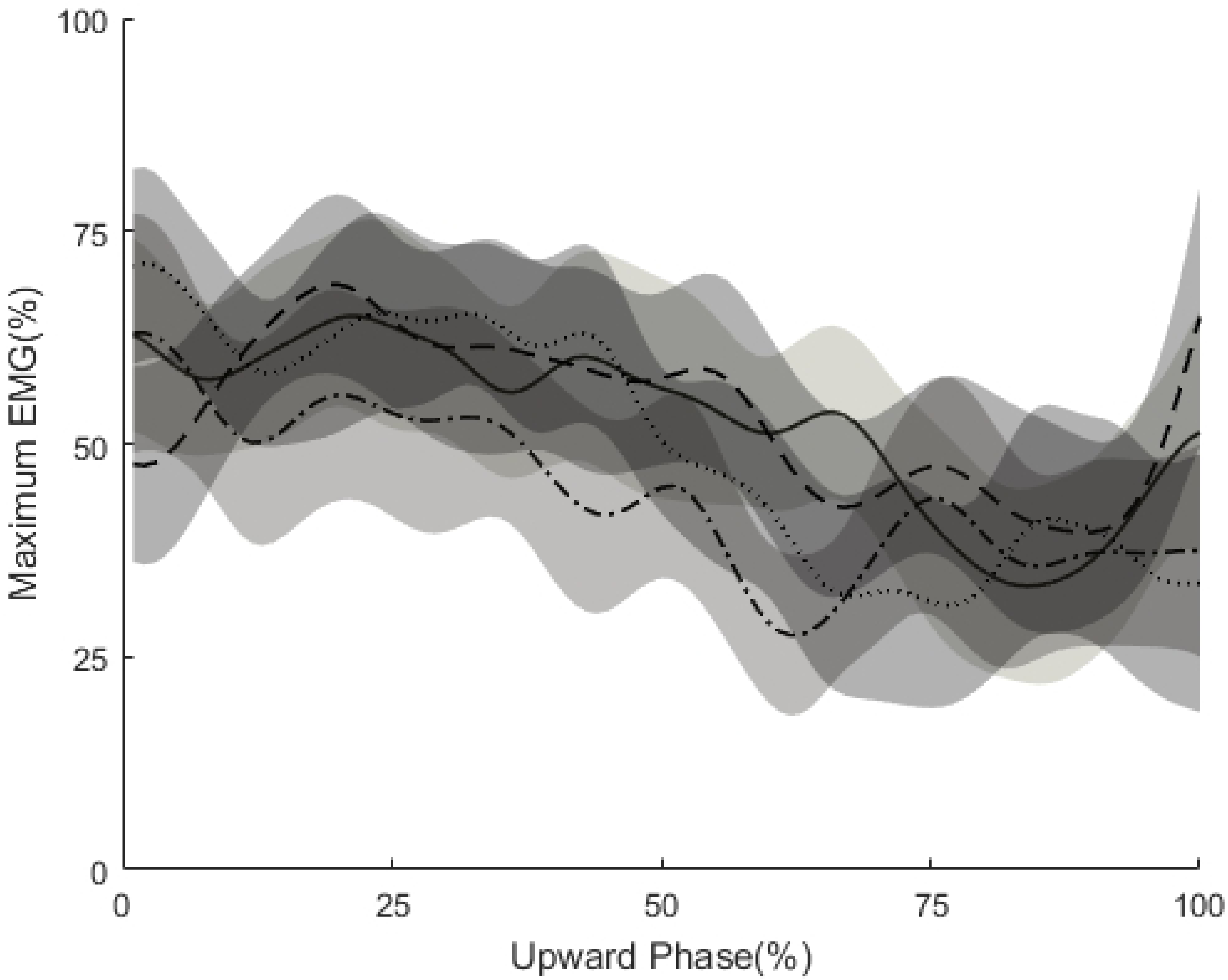
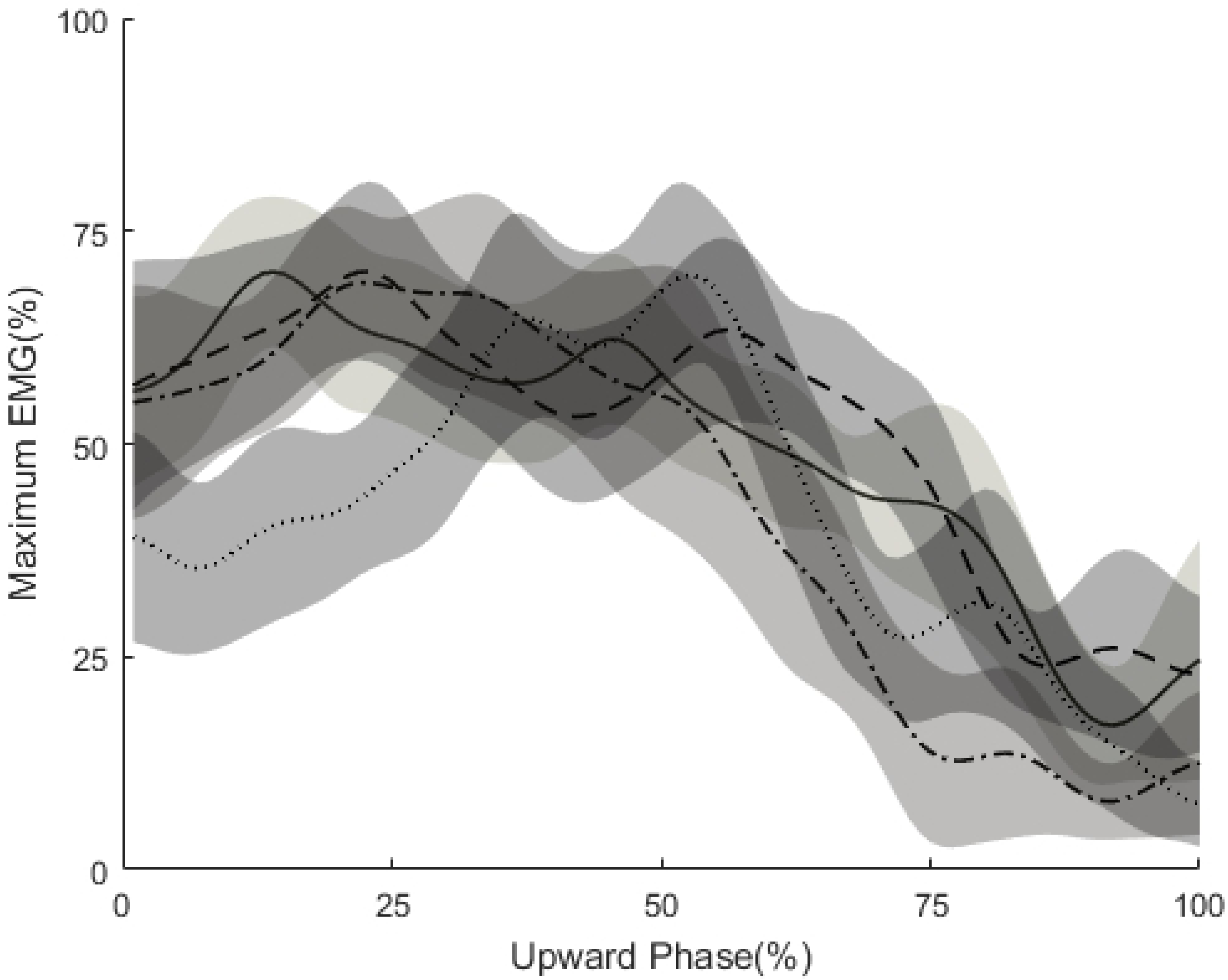
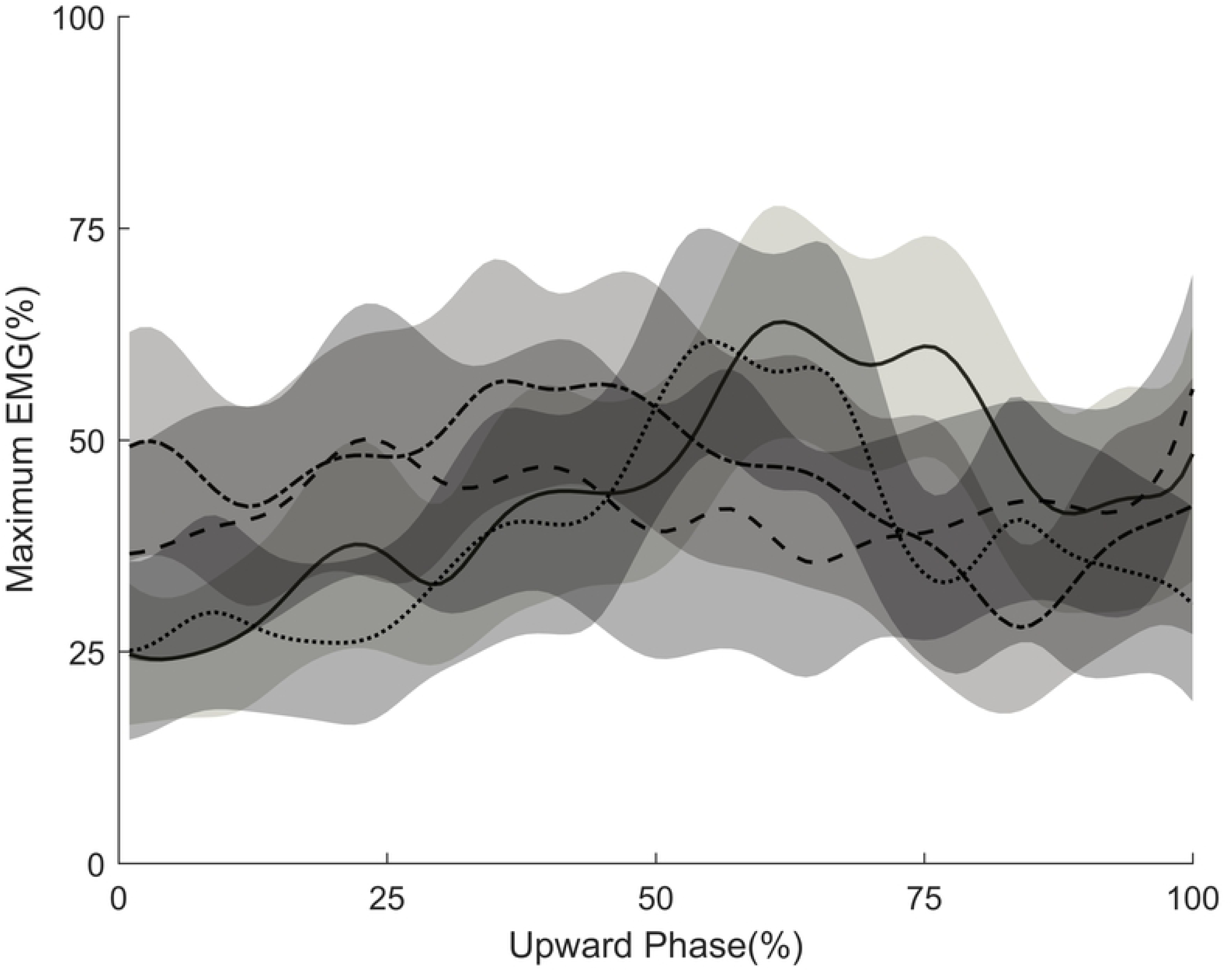
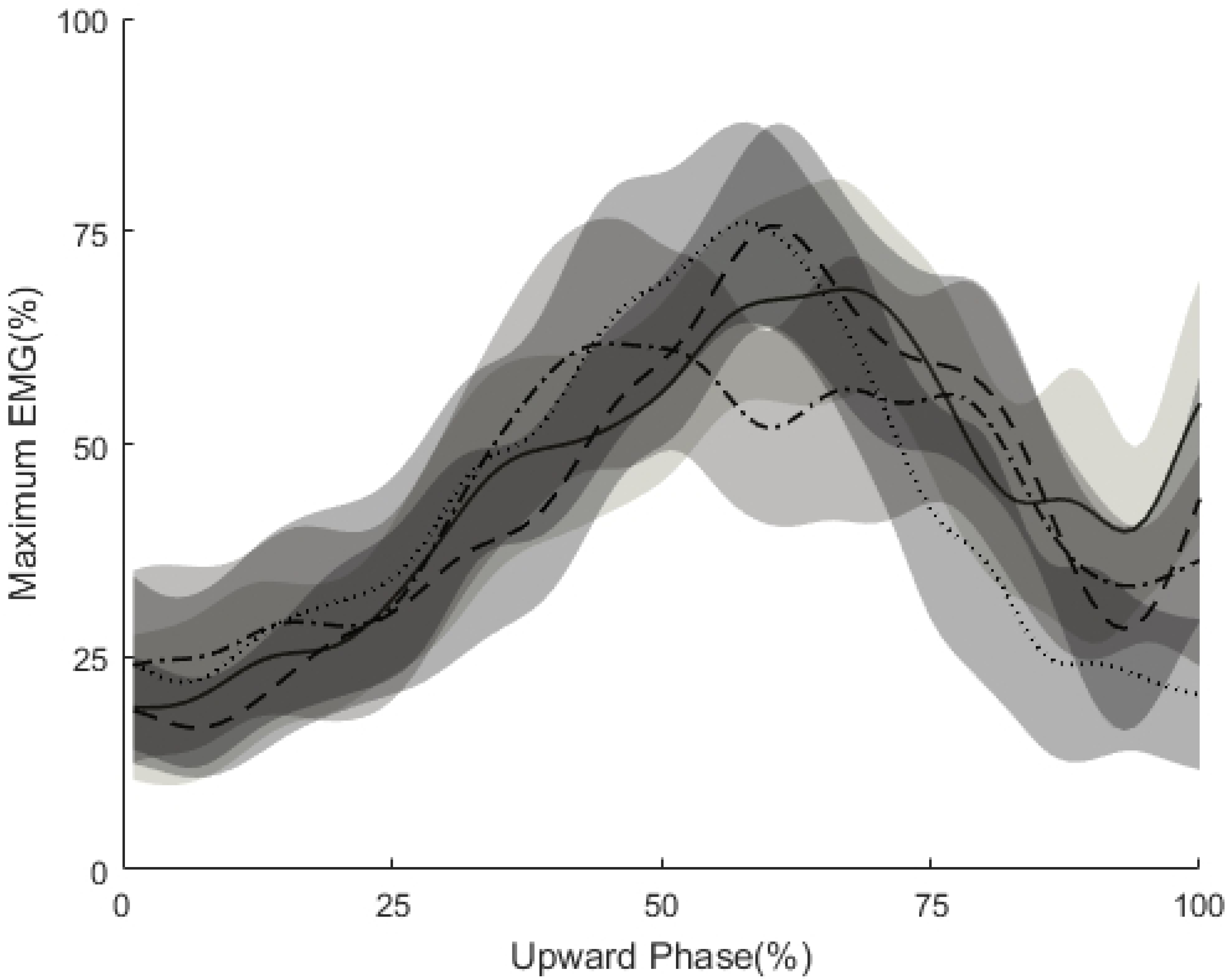
Linear enveloped EMG curves for (A) rectus femoris, (B) vastus medialis, (C) gluteus maximus, (D) biceps femoris, and (E) medial gastrocnemius as a percentage of the maximum signal attained during the upward phase of the jump squat, jump, back jerk and front jerk. Curves are ensemble averages for all participants, while the shaded bands represent the 95% confidence interval.

While normalization of each EMG values facilitates a statistical comparison, it is necessary to compare the peak EMG values for every muscle between exercises. The rectus femoris was activated to a significantly greater level in vertical jump than front jerk (F(1.241, 23.582) = 3.051, P < .1, η^2^_p_ = .138) (Fig6). The vastus medialis exhibited the same features although the significant greater level was in jump squat compared to back jerk and no significant differences between other exercises (F(2.064, 39.217) = 3.580, P < .05, η^2^_p_ = .036) (Fig6). The gluteus maximus was activated to a significantly greater level in front jerk than vertical jump (F(3, 57) = 5.838, P < .005, η^2^_p_ = .235)(Fig6).Although not statistically significant, biceps femoris and medial gastrocnemius showed the same patterns as in two quadriceps muscles (Fig6).

**Fig 6.**
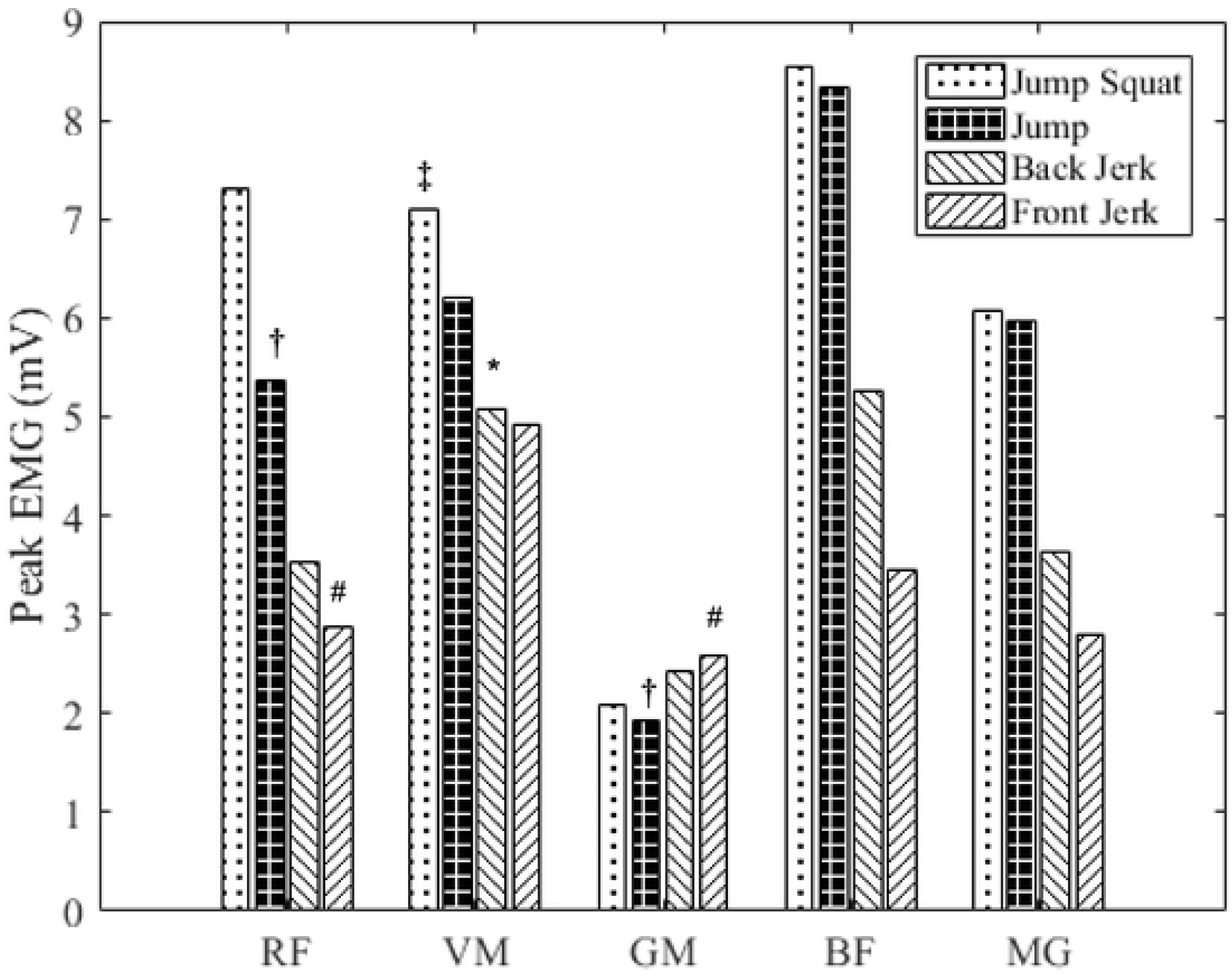
Peak EMG values during the jump squat, jump, back jerk and front jerk for each of the five muscles tested. #indicates significantly different from vertical jump (P ≤ .05). *indicates significantly different from jump squat (P ≤ .05). †indicates significantly different from front jerk (P ≤ .05). ‡indicates significantly different from back jerk (P ≤ .05).

## Discussion

To knowledge of the author, this is the first research to compare the EMG character of WOPDs, jump squat and vertical jump. Similar characters were exhibited between jump squat and vertical jump, which was consistent with previous research that there were no significant differences of peak EMG between vertical jump and jump squat in vastus medialis, rectus femoris, biceps femoris, gluteus medius and medial gastrocnemius [29]. Instead of gluteus medius, current study assessed gluteus maximus and still no significant difference existed between jump squat and vertical jump. However, significant differences in peak EMG were showed between WOPDs and vertical jump. Jerk exhibited a significantly lower value in rectus femoris and a significantly higher value in gluteus maximus than vertical jump. Back jerk exhibited a significantly lower value than jump squat in vastus medialis. The timing of muscles reached their peak value showed that jerk was significantly different than vertical jump in rectus femoris and gluteus maximus. Overall, jerk and back jerk exhibited the same features that lower peak value and earlier timing of peak value reached in muscles around knee and ankle joints which are vastus medialis, rectus femoris and medial gastrocnemius, besides they also exhibited greater peak value and later timing of peak value reached in gluteus maximus, although some of them were not to a significant level in statistics.

The kinematics features were conformed to the EMG results as well. The vector coding technique was used to examine the joint coupling pattern of the exercises for a more integral comprehension of WOPDs and jumping movements in kinematics. Since a value greater than 45° suggested more distal motion relative to proximal motion and vice versa. All the lines of HK and KA indicated that the extensions of hip, knee and ankle of all movements were activated from proximal to distal. This was the reason that weightlifting derivatives and jumping movements were considered similar in lower-limb kinematics [26]. If the most similar of movement is the first and foremost factor in exercise selection, the details have to be compared in further depth. The triple extensions of vertical jump and jump squat showed a clear shift in dominance from hip to knee then ankle, although ankle flexion was on-going at the beginning of knee extension. This allows transfer of energy in a more optimal sequence between joints. Accordingly, jumps squat was the most similar movement with jump in both HK and KA coupling patterns.

Both jerk and back jerk were significantly different with vertical jump in HK coupling pattern (Fig1A). It exhibited a more knee dominant motion throughout the upward phase. From 60% of upward phase, it was obvious in the KA coupling angle (Fig1B) that jerk and back jerk was overlapped and motions were more ankle dominant than two jump conditions. Only jump squat was similar to vertical jump in both HK and KA coupling patterns. Other researchers also reported that the loaded jump squat was more correlated to jumping and sprinting abilities than the push press [20, 21]. It may be explained by different strategies were used in jumping and jerking movements. Jump was dominated by knee and hip in a more balance recruitment, whereas jerk represented a more knee dominant activity [18]. After the comparison of the joint moment and impulse between jerk, jump squat and a countermovement jump (CMJ), Cushion found that the greatest mechanical similarity with CMJ was observed at lower relative loads of the jerk and jump squat [19]. The mechanism behind is that subjects would alter the strategy in which they carry out the jumping and jerking as load increased such that the resemblance to CMJ characteristics is altered [19]. In current study the 40% 1RM squat was used as an absolute load to each exercise, which is much higher than the load of 40% 1RM jerk [30]. Thus, the dissimilarity between both jerk conditions to vertical jump was reinforced by the relative higher load in jerk trials.

The other possible explanation was that the center of gravity of the system (bar plus body mass) and that of the barbell do not move in parallel [31–34] during weightlifting. Such as clean, jerk, snatch and other variations, these movements emphasizes on developing high levels of muscular power and to apply that power to the bar in a brief period[35]. Accordingly, the focus of a weightlifter is moving an external object as quick as possible because the success of weightlifting is determined by the power transferred to the barbell [34, 36]. The difference is that jumping movements are to overcome the inertia of center of the system. It may also explain the earlier timings of jerk and back jerk reached in peak kinetics values and some of the peak EMG values. In comparison with both jumping movements, all kinetic variables of both jerk conditions, such as peak force, peak power and peak RFD, showed the earlier timings reached in current study. They were required to transfer the power to the bar quicker than jumping conditions.

In the past, studies suggested similarities in kinetic parameters with push press, power clean and jump squat. Lake et al. [30] found the push press enabling application of significantly greater power with less mechanical cost than the jump squat equivalent and suggested that push press could provide a time-efficient combination and effective stimulus to the entire kinetic chain. On the other hand, Comfort et al. [26] indicated that the push press, squat jump and mid-thigh power clean could be replace by each other without affecting the training of peak power and rapid force production when the load was at 50-70% 1RM. Moreover, Comfort et al. [37] reported under the load of 60% 1RM power clean, there were no differences in peak force, peak power or rate of force development (RFD) with the push press, squat jump and mid-thigh power clean, concluding that all these exercises could be used interchangeably for power and force development with moderate load. The difference of push press and jerking exercises was the catching phase after impulsive triple extension. The jerking exercises performed a drop under the bar by athletes lowering the center of mass to catch the bar in the overhead position, which allows a heavier load lifted than push press. Considering jerking exercises share the same propulsion phase with push press, it was reasonable to refer previous researches including push press.

In the contrary, key findings of this study in kinetics were that: (a) Peak force was similar between back jerk and jump squat, while both conditions were significantly greater in than jerk; (b) Peak RFD was different that although in the upward phase there was no significant differences between jump squat and both jerks, considering all four exercises were countermovement and the purpose of the present study was to find which exercise was better in improving vertical jump, therefore the conclusion was that jump squat and back jerk were significantly greater than jerk including eccentric phase. (c) Peak power was significantly higher in jump squat than both jerks while back jerk was significantly higher than jerk. In summary, jump squat elicited greater results than jerk in almost all kinetics parameters and back jerk was more similar to jump squat in kinetics.

The different results was attributed to two reasons. Firstly, previous studies used the relative load for each exercise. In this study, 40% 1RM squat was used as the same load for all three resisted lifts instead of relative load, which was indicated as the optimal load for peak power of jump squat [38]. With previous study indicated the peak power of jerk and back jerk occurred at 80-90% 1RM of themselves, thus the load of 40% 1RM squat may not be the optimal load for WOPDs [23]. If different loads were used by each movements, it may cause a disturbing effect for kinematics comparison between WOPDs (jerk and back jerk) and traditional ballistic exercises (loaded jump squat) as suggested in previous studies [19]. Secondly, the force plate calculates power from the ground reaction force and velocity of the center-of-mass as integrated from the acceleration-time curve [39] and as we can observed in Fig 4B there was a greater peak force and longer duration of jump squat which may cause the greater value of power than both jerk conditions.

The second purpose of this study was to explore the biomechanical differences resulted from initial position of the bar in jerking movements. Only one research conducted a comparison of power output across a spectrum of loads (30-90%) with jerk and back jerk. It indicated that relative intensity had a significant impact on peak power with jerk and back jerk and also reported that the back jerk elicited greater power output than the jerk for all the loads assessed [23]. It was supported by our findings that back jerk (2223N and 3767W) was greater than jerk (2061N and 3364W) in peak force and peak power respectively. Researches that have studied the kinetic parameters of the jerk performance have reported very high loads lifted as well as great values of power outputs have been developed [14,41,42]. Some previous findings have shown a lower peak power than this study: Flores et al. [23] and Lake et al. [40] reported values of 3103 W and3046 W during the jerk respectively. As well, Garhammer [41, 42] reported mean power was from 2690–4321W and peak power was from 2500-6953W during the jerk, which include our findings in the range.

In rate of force development jerk (7702 N·s^−1^) was significantly greater than back jerk (5972 N·s^−1^) in upward phase, but the opposite in downward phase that back jerk (15965 N·s^−1^) was significantly greater than jerk (12280 N·s^−1^). In summary, findings from this study were that no significant difference elicited by jerk and back jerk in joint coordination and EMG characters and back jerk surpass jerk in almost all kinetics parameters. As it was previously suggested, the reason may be behind the nature of the movement. The chin of the lifter was on the trajectory of the barbell upward in jerk. And back jerk with no obstacles, which elicited an easier technical pattern than jerk, allowed development of higher force and higher velocity [31–33]. In addition, jerk from the chest required a sound shoulder flexibility to lie the bar on the anterior deltoid. In practical experiences, this flexibility was scarce in large sport populations meanwhile back jerk was more applicable with individuals as long as no shoulder dysfunctions suffered.

An interesting result of this study was that vertical jump was significantly greater in peak power and peak RFD among all four tested movements. It was suggested that the load of maximizing power output between the bar, body, and system was 80%, 0%, and 0% of 1-RM respectively. That means jump without load elicited a greater power output than any load used [34]. Moreover, MacKenzie et al. compared jump squat, power clean and vertical jump and reported no significant differences in peak power and peak RFD between jump squat and vertical jump [29]. Due to half of his subjects were females, it was obvious by scrutinizing his diagram that the male counterpart was obviously higher in vertical jump than jump squat. Furthermore, the instructions of countermovement jump condition was different as well. The arm swing of vertical jump in current study was free to swing during all downward and upward phases, during which in MacKenzie’s research, arms were still at their sides during the downward phase and only free to swing during the upward phase of the movement [29].

In addition, the level of the subject in experience and proficiency had an influence on the percentage of peak strength at which the greatest power is developed either upward or downward, as suggested previously [10]. In such a way, it would be less clear with the impact from the strength level of the subjects. Subjects of the current study were younger sprinters or jumpers in professional level, instead of either professional weightlifters or some resistance or recreational trained man, which may explain the higher performance in vertical jump than lifts and also has more reference to practitioners and coaches. Although modalities such as Olympic weightlifting derivatives and lower-limb ballistic exercises were often employed to improve vertical jump ability, the result from this study suggested that the vertical jump is a valuable training exercise in its own right. Vertical jump itself is an irreplaceable training tool in improving the jumping ability. Further research could consider whether loaded jump with arm swing such as wearing a weight vet or using vertimax would be a better training exercise for improving jumping height.

## Conclusions

Current study found that jump squat was more similar to vertical jump than WOPDs in kinematics and EMG character. Jump squat and back jerk were better than jerk in force and power production during concentric phase and in RFD during eccentric phase. Back jerk was more similar to jump squat in kinetics than jerk. In view of subjects of the current study was not professional weightlifters or some resistance or recreational trained man like most previous researches used. Therefore, in the sport populations such as young runners and jumpers, jump squat was more recommended for improving jumping ability than WOPDs. Back jerk was more recommended than jerk for athletes who targeted the ability to develop rapid force production and power development. In addition, vertical jump is a good training exercise by itself.

## Acknowledgments

The author express his appreciation to the subjects for their participation in this study as well as staffs and students from Scientific Research Center and Sport Biomechanical Department of Beijing Sports University who carried out the experiment and offered support.

## Supporting information

**S1 Fig1. Upward phase of the (A) hip-knee coupling angle and (B) knee-ankle coupling angle during the upward phase of the jump squat, jump, back jerk and front jerk.** Curves are ensemble average for all participants, while the shaded bands represent the 95% confidence interval.

**S2 Fig2. (A) Peak vertical ground reaction force, (B) peak power generated and (C) peak rate of force development (RFD) during the upward phase of the jump squat, jump, back jerk and front jerk.** #indicates significantly different from vertical jump (P ≤ .05). *indicates significantly different from jump squat (P ≤ .05). †indicates significantly different from front jerk (P ≤ .05). ‡indicates significantly different from back jerk (P ≤ .05)

**S3 Fig3. (A) Vertical ground reaction forces and (B) power as a percentage of the maximum values attained during the upward phase of the jump squat, jump, back jerk and front jerk.** Curves are ensemble averages for all participants, while the shaded bands represent the 95% confidence interval.

**S4 Fig4 (A) Typical vertical ground reaction force and (B) power profiles, for Participant 7, for the jump squat, jump, back jerk and front jerk.** Curves start at the initiation of the downward phase, while the shaded portions indicate the upward phase. Curves end when the weight of the participant, or participant and barbell, exceeded the magnitude of the vertical ground reaction force.

**S5 Fig5. Linear enveloped EMG curves for (A) rectus femoris, (B) vastus medialis, (C) gluteus maximus, (D) biceps femoris, and (E) medial gastrocnemius as a percentage of the maximum signal attained during the upward phase of the jump squat, jump, back jerk and front jerk.** Curves are ensemble averages for all participants, while the shaded bands represent the 95% confidence interval.

**S6 Fig6 Peak EMG values during the jump squat, jump, back jerk and front jerk for each of the five muscles tested**. #indicates significantly different from vertical jump (P ≤ .05). *indicates significantly different from jump squat (P ≤ .05). †indicates significantly different from front jerk (P ≤ .05). ‡indicates significantly different from back jerk (P ≤ .05).

